# Nitrogen assimilation plays a role in balancing the chloroplastic glutathione redox state under high-light conditions

**DOI:** 10.1101/2023.03.27.534362

**Authors:** Gal Gilad, Omer Sapir, Matanel Hipsch, Daniel Waiger, Julius Ben-Ari, Nardy Lampl, Shilo Rosenwasser

## Abstract

Nitrate reduction and subsequent ammonium assimilation require reducing equivalents directly produced by the photosynthetic electron transport chain. Therefore, it has been suggested that nitrate assimilation provides a valuable sink for excess electrons under high-light (HL) conditions, which protects the photosynthetic apparatus from excessive harmful reactive oxygen species. This work experimentally tested this hypothesis by monitoring photosynthetic efficiency and the chloroplastic glutathione redox state (chl-*E*_*GSH*_) of plant lines with mutated glutamine synthetase 2 (GS2) and ferredoxin-dependent glutamate synthase 1 (GOGAT1), two key enzymes of the nitrogen assimilation pathway. Unlike wild-type (WT) plants, mutant lines incorporated significantly less isotopically-labeled nitrate into amino acids, demonstrating impaired nitrogen assimilation. When nitrate assimilation was compromised, photosystem II (PSII) proved more vulnerable to photodamage, as shown by the low PSII quantum yields recorded in the mutant lines. High temporal resolution monitoring of the redox state of chloroplast-targeted reduction-oxidation sensitive green fluorescent protein 2 (chl-roGFP2), expressed in the background of the mutant lines, enabled assessment of the effect of the nitrate assimilation pathway on the chl-*E*_*GSH*_. Remarkably, while oxidation followed by reduction of chl-roGFP2 was detected in WT plants in response to HL, oxidation values were stable in the mutant lines, suggesting that the relaxation of chl-E_GSH_ after HL-induced oxidation is achieved by diverting excess electrons to the nitrogen assimilation pathway. Together, these findings indicate that the nitrogen assimilation pathway serves as a sustainable energy dissipation route, ensuring efficient photosynthetic activity and fine-tuning redox metabolism under light-saturated conditions.

## Introduction

Growing under highly variable light conditions, plants must constantly sense and adjust to daytime photon fluxes. Under high light conditions (HL), NADPH and ATP production in the light reactions might exceed their consumption in the downstream Calvin-Benson cycle (CBC). Efficient coupling between energy generation and consumption is required to avoid damage to photosystems and over-reduction of the chloroplastic electron transport, maintaining photosynthesis-derived reactive oxygen species (ROS) under harmless low levels. Several energy dissipation mechanisms, including non-photochemical chlorophyll fluorescence quenching (NPQ) (Biehler and Fock, 1996; Asada, 1999; Ort and Baker, 2002; Miyake, 2010), the water-water cycle (WWC) (Asada, 1999; Mehler, 1951) and plastid terminal oxidase (PTOX) activity (Nawrocki et al., 2015; Kambakam et al., 2016; Saroussi et al., 2019; Rog et al., 2022), have evolved in photosynthetic organisms to quench excessive light energy and achieve homeostasis and optimal photosynthetic performance under HL. Compared with the linear electron transport in which electrons derived from water splitting in PSII reduce NADP+, the activity of energy dissipation mechanisms leads to a net loss of photonic energy or reducing equivalents, protecting from excess reductive activity and helping in adjusting ATP/NADPH ratio. An additional way to protect the photosynthetic apparatus from excess illumination while conserving energy is to channel energetic electrons into reduced carbon products such as starch and lipids (Treves et al., 2016; Saroussi et al., 2019).

Besides carbon fixation, photosynthesis generates ATP and reducing power that is used to assimilate inorganic nutrients into organic substances, with nitrate assimilation being the most energy-consuming process. Compared with CO_2_ reduction, nitrate assimilation requires more electrons and less ATP, thus playing a role in balancing ATP/NADPH (Noctor and Foyer, 1998). However, the direct role of nitrogen assimilation in the alleviation of HL-induced photodamage and maintaining chloroplast redox homeostasis is not fully resolved.

In most environments, nitrate (NO_3_^−^) is the primary nitrogen form available (Miller et al., 2007). Its assimilation occurs in the roots and shoots of vascular plants, with the shoot being the predominant site in most cases (Scheurwater et al., 2002; Xu et al., 2012). The nitrate assimilation pathway consists of several key enzymes and is spatially separated between the cytoplasm and plastids/chloroplasts. Nitrate reductase (NR) catalyzes nitrate reduction in the cytosol, forming nitrite, a highly reactive nitrogen compound. Nitrite is transported from the cytosol to the chloroplast, where it is reduced to ammonium via nitrite reductase (NiR) activity. The ATP-dependent incorporation of ammonium into amino acids is catalyzed by the glutamine synthetase (GS)/glutamate synthase (GOGAT) cycle (Lea and Miflin, 1974), in which GS catalyzes the production of glutamine (Gln) from ammonium and glutamate (Glu). The reductive transfer of the amide group from Gln to α-ketoglutarate, which produces two Glu molecules, is catalyzed by GOGAT. Glu and Gln serve as N-donors for the production of various N-containing compounds, such as the biosynthesis of amino acids and nucleic acids (Coruzzi, 2003).

Two types of GS were found in plants; GS1 and GS2 (Lea and Miflin, 2003). In Arabidopsis, GS1 type is encoded by five genes (GLN1;1-5) and is active in the cytosol of roots and leaf cells. GS2 is encoded by a single gene (GLN2) and is located in plastids/chloroplasts. It has been suggested that GS2 plays a central role in the nitrate assimilation pathways as well as in the reassimilation of NH_4_^+^ released during photorespiration (Bernard and Habash, 2009; Ferreira et al., 2019; Marino et al., 2022). Indeed, barley, legume Lotus japonicus and tomato mutants lacking GS2 failed to thrive and showed severe stress symptoms under normal air conditions while growing normally under non-photorespiration conditions (i.e., O_2_-depleted or CO_2_-enriched air) (Blackwell et al., 1987; Wallsgrove et al., 1987; Orea et al., 2002). In contrast, the recently characterized Arabidopsis GS2 mutants had no severe phenotypes, demonstrating that GS2 activity is not critical for plant survival and questioning its central role in primary N metabolism (Ferreira et al., 2019; Hachiya et al., 2021; Lee et al., 2022; Marino et al., 2022).

NADH-dependent and ferredoxin (Fd)-dependent GOGAT enzymes have been identified in plants, with the latter being unique to photosynthetic organisms and responsible for most GOGAT activity in leaves (Somerville and Ogren, 1980; Suzuki and Rothstein, 1997). In the Arabidopsis genome, two genes encoding Fd-GOGAT were found: GLU1 and GLU2, with GLU1 playing an active role in re-assimilating photorespiratory NH_4_^+^ as well as in nitrogen assimilation in shoots, whereas GLU2 is involved in root nitrogen assimilation (Coschigano et al., 1998).

Nitrate reduction and subsequent ammonium assimilation require a considerable amount of reducing equivalents in the form of Fd or NADPH, which, in leaves, are directly supplied by the photosynthetic electron transport chain (Stitt et al., 2002; Foyer et al., 2011; Buchanan et al., 2015). Depending on the environmental conditions and nitrate availability, nitrate reduction can consume 5-25% of reducing equivalents generated in chloroplasts, rendering it the second largest sink for low potential electrons generated by the photosynthetic electron transport chain (Bloom et al., 1989; Edwards and Baker, 1993; Stitt et al., 2002). Accordingly, nitrate assimilation is considered the primary factor contributing to the gap between electron requirements for oxygen evolution and CO_2_ assimilation (de la Torre et al., 1991; Edwards and Baker, 1993; Noctor and Foyer, 1998).

ROS detoxification via the ascorbate–glutathione cycle involves drawing of electrons from the glutathione (GSH) pool (Foyer and Noctor, 2011; Awad et al., 2015)). The resultant GSH disulfide (GSSG) is reduced by the electron carrier NADPH in a reaction catalyzed via GSH reductase (GR). Thus, changes in the balance between ROS and reducing power production can be reflected in the GSH redox potential (*E*_*GSH*_), which is dependent on the [GSH]/[GSSG] ratio (Foyer and Noctor, 2011; Rahantaniaina et al., 2013). Recently the role of chloroplastic *E*_*GSH*_ in maintaining efficient photosynthesis has been demonstrated using GR knockout lines in the model moss *Physcomitrella patens* (Müller-Schüssele et al., 2020). High spatiotemporal resolution monitoring of *E*_*GSH*_ can be achieved using redox-sensitive green fluorescent proteins (roGFPs) (Meyer et al., 2007; Meyer, 2008; Meyer and Dick, 2010), which carry two engineered cysteine residues that form an intramolecular disulfide bridge which impacts its fluorescence characteristics (Dooley et al., 2004; Hanson et al., 2004). Recently, daily patterns in chloroplast-specific *E*_*GSH*_ under normal and HL conditions and the effect of NPQ and cyclic electron flow pathways in shaping these patterns were resolved using chloroplast-targeted roGFP2 (Haber et al., 2021). However, the possible role of nitrate assimilation in preventing over-reduction and ROS generation under HL conditions, thereby balancing the chl-*E*_*GSH*_, has not been investigated.

The present work aimed to examine the possible role of the nitrogen assimilation pathway as an alternative reducing power sink under HL by examining the flux from nitrate into glutamine and glutamate, PSII efficiencies and chl-E_GSH_ dynamics in plant mutated in GS2 and GOGAT1. We demonstrated the induction of nitrate assimilation pathways under HL and the impairment of de-novo nitrogen assimilation in the examined mutant lines. We also showed that the decrease in nitrate assimilation rates observed in the mutant lines is associated with a lower PSII efficiency and abnormal daily chl-*E*_*GSH*_ dynamics.

## Results

### Growth phenotype of GS2 and GOGAT1 mutant lines

To assess the essentiality of nitrate assimilation as an energy dissipation route, *gln2* (Ferreira et al., 2019) and *glu1* (Somerville and Ogren, 1980) mutant lines, defective in GS2 and GOGAT1, respectively (Fig. 1A) were studied. Immunoblot analysis confirmed a significant reduction in GS2 and GOGAT1 levels in *gln2* and *glu1* lines, respectively, compared to WT (Fig. 1B). The appearance of weak bands in protein extractions derived from the mutant lines may reflect residual expression levels or coincidental non-specific binding of the antibody. No severe phenotype was observed in *gln2* mutant lines despite being slightly smaller than WT. For example, *gln2* leaf area was ∼70% of that of WT plants (Fig. 1D-F). These results are in agreement with recent reports which showed that Arabidopsis *gln2* knockout mutants are viable under photorespiratory conditions (Lee et al., 2022), but lie in contrast to the severe phenotype of GS2 mutants in barley and *L. japonicus* (Blackwell et al., 1987; Orea et al., 2002). The *glu1* plants showed a chlorotic phenotype and were small, with a five-fold reduction in leaf area compared to WT plants. These results are in agreement with previous observations of the typical photorespiratory phenotype of *glu1* plants (Somerville and Ogren, 1980; Coschigano et al., 1998)

**Figure 1:**
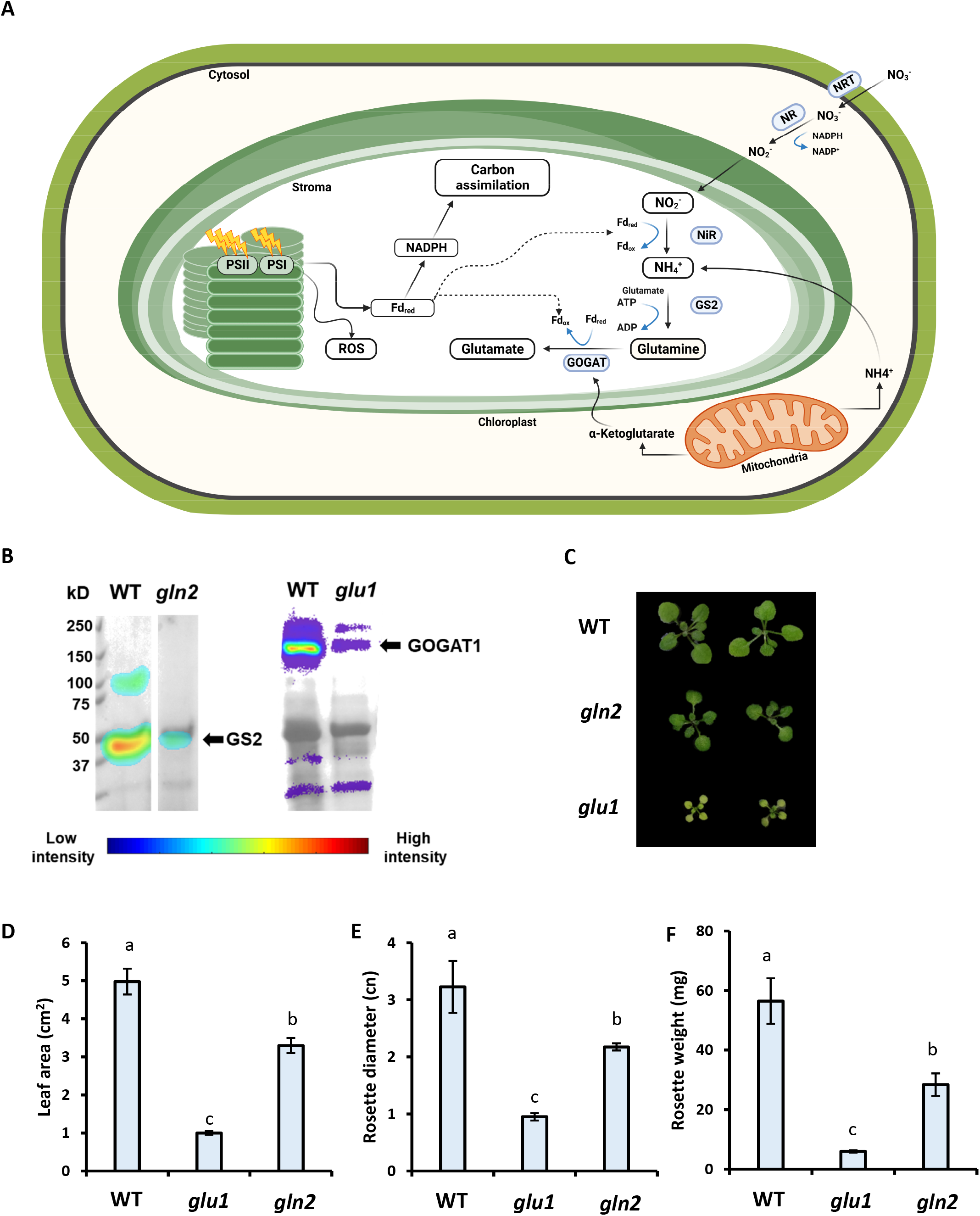
The nitrogen assimilation pathways and mutant lines used in this study. (A) The nitrogen assimilation pathway and photosynthesis are interconnected. Fd serves as an electron carrier for electrons generated by photosystem I and consumed in the nitrogen and carbon assimilation pathways. In the nitrogen assimilation pathway, Fd donates an electron to mediate the reduction of nitrite to ammonium and the conversion of Gln to Glu. Created with BioRender.com. (B) Western blotting using anti-GS2 and GOAGT1 antibodies of *gln2* and *glu1* plant lines. WT was used as a positive control. (C) Representative photographs of *gln2* and *glu1* mutant lines. (D-F) Characterization of the *gln2* and *glu1* at three weeks old. Lead area (D), average rosette diameter (E), and weight (F) are presented. Data represent the average ± SE (n=4).

### The primary nitrate assimilation pathway involves GS2 and GOGAT1 activity and is enhanced under HL

To directly explore the *in vivo* activity of the nitrogen assimilation pathway, the rate of nitrogen assimilation was evaluated by following the transfer of isotope-labeled ^15^N from nitrate into Gln and Glu in hydroponically grown plants (Fig. 2A). Gln contains both amide and amine nitrogen groups, while Glu contains one amine.

**Figure 2:**
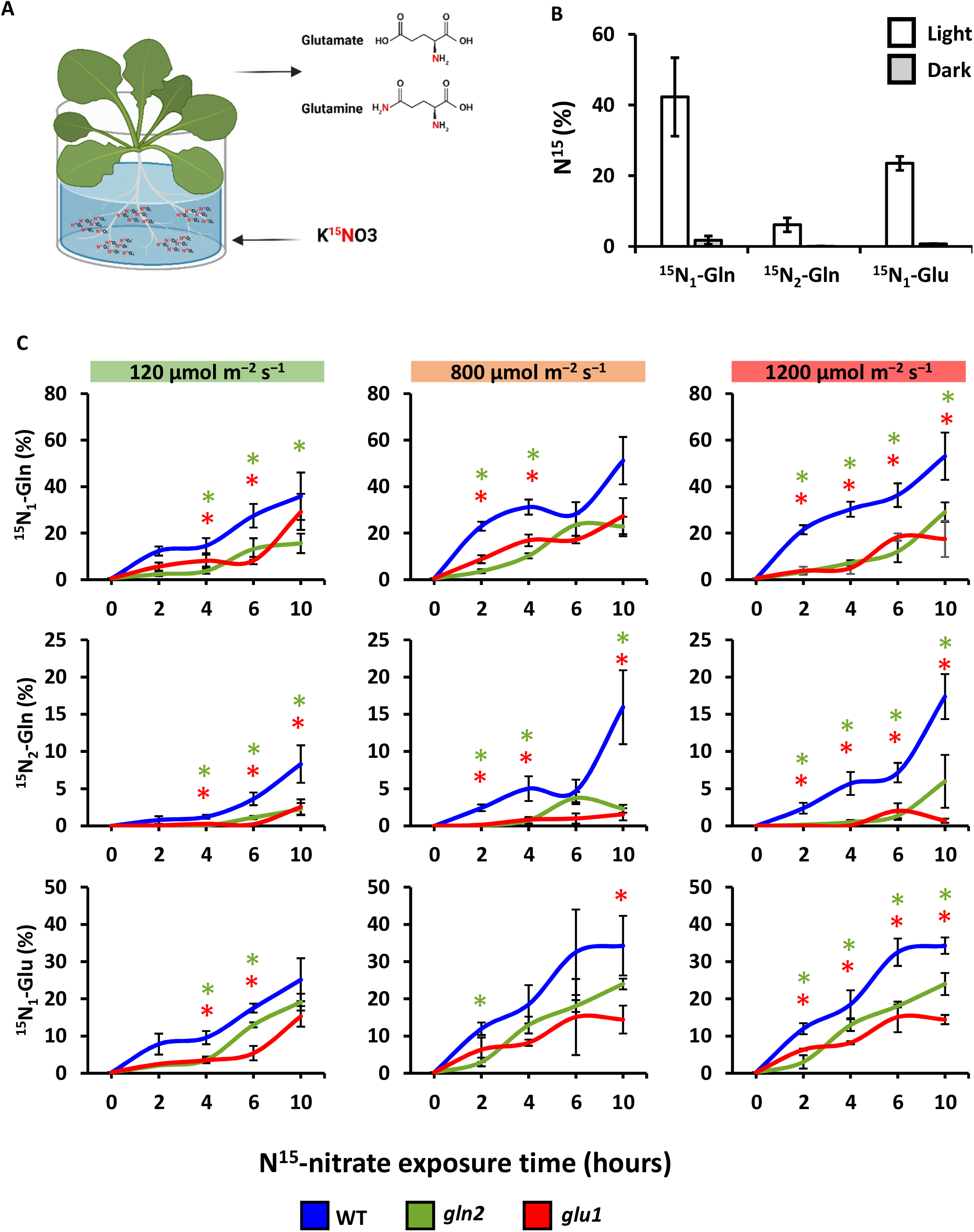
Nitrate assimilation rates in *gln2* and *glu* are lower than in WT. (A) ^15^N-nitrate assimilation into glutamine and glutamic acid was assessed in hydroponically-grown plants. Created with BioRender.com. (B) The relative abundance of ^15^N-labeled Gln and Glu amino acids in WT Arabidopsis plants grown under normal growth conditions (120μmol m^−2^ s^−1^, 16 h light/8 h dark) or kept in the dark for 24 h, is shown. Labeled amino acids were detected by LC-MS. ^15^N_1_-Gln and ^15^N_2_-Gln refer to Gln labeled with one or two ^15^N atoms, respectively. Values are presented as means of 4 plants ± SE. (C) Nitrogen assimilation rates in WT, *gln2* and *glu1* lines exposed to different light intensities. The data are presented as an average of 3 plants ± SE. Green and red asterisks indicate significant differences (p ≤ 0.05) between WT and *gln2* or *glu*, respectively.

Accordingly, labeled Glu was detected by a 1Da mass shift while labeled Gln was identified as either a 1Da or 2Da mass shift, indicated as ^15^N_1_-Gln and ^15^N_2_-Gln, respectively. The analysis was carried out using high-resolution mass spectrometry, allowing the resolving of naturally occurring ^13^C isotopes from experimentally introduced ^15^N isotopes (Figures S1-S3). Isotopically-labeled ^15^N, derived from ^15^NO_3_^-^ was efficiently incorporated into Gln and Glu in WT plants maintained for 24 h under normal growth conditions (NL, 120 μmol m^−2^ s^−1^, 16 h light/8h dark), with ∼23.5% of the total Glu marked with ^15^N and ∼42.3% and ∼6.1% of the total Gln marked with ^15^N_1_ and ^15^N_2_, respectively (Fig. 2b). The structure of the nitrogen assimilation pathway can explain this hierarchy in the rate of accumulation of the labeled amino acids. As shown in Fig. 1A, Gln is the first amino acid assimilated with labeled nitrogen via GS activity, explaining its fastest labeling rate. Then, in a GOGAT-catalyzed reaction, the amide group of Gln is transferred to a 2-oxoglutarate molecule, yielding two glutamates. The lower abundance of ^15^N-Glu compared to ^15^N_1_-Gln is likely the result of diversion of some of ^15^N_1_-Gln to amino acid or protein biosynthesis instead of to ^15^N-Glu. As predicted, the relative abundance of ^15^N_2_-Gln was lower than ^15^N_1_-Gln, as it was generated through GS2 activity with ^15^N-Glu serving as a substrate. Minor ^15^N incorporation into Glu and Gln was observed in plants maintained for 24 h in the dark, with the percentage of labeled amino acids being 0.73%, 1.75% and 0.04% for ^15^N-Glu, ^15^N_1_-Gln and ^15^N_2_-Gln, respectively. In comparison, values less than 0.006 were recorded in leaf samples not exposed to ^15^N, representing naturally occurring isotopes. These results clearly demonstrate the light dependency of the nitrogen assimilation pathway (Fig. 2B).

To determine whether GS2 and GOGAT1 activities are required for the *de-novo* nitrate assimilation pathway and how light intensities affect this process, WT, *gln2* and *glu1* plants, grown under NL, were treated with ^15^NO_3_^-^ and immediately exposed to 800 (ML) or 1200μmol m^−2^ s^−1^ (HL) for 10 h or left under NL as control (Fig. 2C). An increase in the relative abundance of labeled Gln and Glu was detected during the experiments in all tested lines, demonstrating their ability to acquire and assimilate the labeled nitrate. Higher incorporation rates were measured under HL compared to NL in WT plants, with, for example, 27.4% and 36.3% of ^15^N_1_-Gln after 6 h under NL and HL, respectively. The most pronounced effect of HL treatment was evident at 4 h and 6 h, (*p* = 0.01, 0.0064, 0.04, for ^15^N_1_-Gln, ^15^N_2_-Gln and ^15^N-Glu, respectively). An increase in the average relative abundance of labeled amino acids was also detected in plants after 4 h and 6 h under ML compared to NL, although the effect of ML was less significant. These results show that high light intensities stimulate the nitrogen assimilation pathway.

Under all three light regimes and throughout the experiment, significantly lower rates of labeled nitrate incorporation were measured in *gln2* and *glu1* compared to WT, demonstrating the impaired nitrogen assimilation pathway in these mutants (Fig. 2c). For example, the relative abundance of ^15^N-Glu after 6 h in NL was 17.5%, 12.9% and 6.2% for WT, *gln2* and *glu1*, respectively. Unlike the light-stimulated nitrate assimilation observed in WT, no significant effect of light intensities on the assimilation rate was detected in *glu1*, while moderate stimulation in *gln2* was observed under ML but not HL. Taken together, the absence of GS2 and GOGAT1 does not entirely abolish nitrogen assimilation but significantly reduces its rate.

### Nitrate assimilation alleviates HL-induced PSII photoinhibition

It was hypothesized that nitrate assimilation under HL prevents damage to PSII by consuming excessive photosynthetic electrons. In such a case, the poor nitrate assimilation in *gln2* and *glu1* was expected to result in a lower electron flux through PSII. To test this hypothesis, ΦPSII was assessed based on chlorophyll fluorescence measurements in WT, *gln2* and *glu1* plants under HL conditions. ΦPSII recorded in darkness, which represents the maximum quantum efficiency of PSII, was lower for *glu1* (ΦPSII =0.73) compared to *gln2* and WT (ΦPSII =0.78), with no significant differences between *gln2* and WT lines (Fig. 3). Severely impaired photosynthetic efficiency was observed in *gln2* and *glu1* plants exposed to HL intensities, reaching ΦPSII values of 0.3-0.4 in plants illuminated at 800 and 1200 µmol m^−2^ s^−1^ (Figure 3 A-B). In comparison, ΦPSII values of approximately 0.6 and 0.5 were recorded for WT plants exposed to the same conditions.

**Figure 3:**
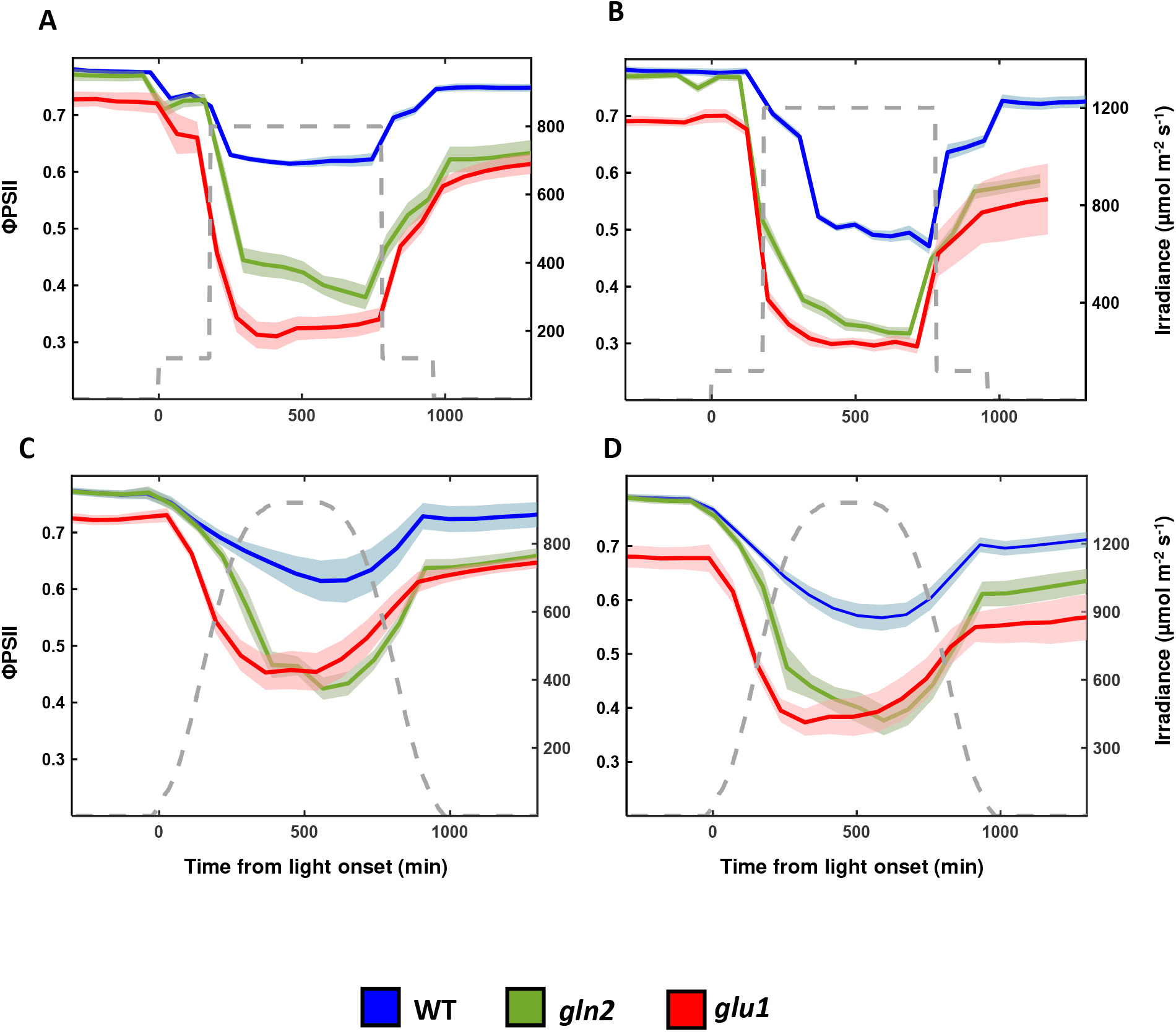
Diurnal changes in ΦPSII under high light conditions in mutants with defects in nitrate assimilation. WT, *gln2* and *glu* lines were subjected to continuous high light (HL) conditions of 800 (A) or 1200 m^−2^ s^−1^ (B) or to gradually increasing light conditions peaking at 800 (C) or 1200 m^−2^ s^−1^ (D). ΦPSII values were derived from chlorophyll fluorescence measurements and are presented as the means of 12 plants, with SE presented as shaded regions.

To mimic physiologically relevant light conditions, plants were exposed to light intensities gradually increasing to a maximum value of either 800 or 1200 µmol photons m^−2^ s^−1^, followed by a gradual decrease in light intensities towards the dark period (Fig. 3 C-D). In general, a decrease in ΦPSII in response to increasing light intensity followed by increasing ΦPSII when the light was dimmed toward the end of the day was found in all three lines, under both light treatments. As expected, in all examined lines, the lowest ΦPSII values were recorded towards the middle of the day, as the light reached its maximum intensity, with the lower values observed in plants exposed to 1200 µmol m^-2^s^-1^ compared to 800 µmol m^-2^s^-1^. Importantly, lower ΦPSII was measured in *gln2* and *glu1* lines compared to WT. For example, mid-day ΦPSII values of 0.44, 0.43 and 0.56 were recorded for *gln2, glu1* and WT, respectively, in response to 800 µmol m^-2^s^-1^ and 0.36, 0.37 and 0.53 were recorded in response to 1200 µmol m^-2^s^-1^. In conclusion, these data demonstrate that impairment of nitrogen assimilation enzymes resulted in lower photosynthetic efficiency, suggesting that the induction of nitrogen assimilation under HL conditions serves as a photoprotective mechanism.

### Gradual relaxation of chloroplastic *E*_*GSH*_ under HL is dependent on the nitrogen assimilation pathway

Considering the role of nitrogen assimilation as an electron outlet, which preserves PSII activity, we hypothesized that it also plays a role in preventing ROS overproduction, thereby shaping chl-*E*_*GSH*_ dynamics under HL. To test this hypothesis, we produced plants expressing the chl-roGFP2 in the background of WT, *gln2* and *glu1* lines. Antibiotic-resistant plants with intense chl-roGFP2 fluorescence levels were selected to monitor the degree of chl-roGFP2 oxidation (Fig. 4A). No phenotypic differences were observed between the original lines and their correspondence lines expressing chl-roGFP2 (Fig. S2). Chloroplast localization of chl-roGFP2 was validated using confocal microscopy, as shown by colocalization of roGFP2 and chlorophyll autofluorescence (Fig. 4B). The probe’s response to redox changes was monitored in the resting state and following the application of hydrogen peroxide (H_2_O_2_) or dithiothreitol (DTT) to induce full probe oxidation and reduction, respectively (Fig. 4C). An increase from resting state 400/488nm fluorescence ratio was observed in H_2_O_2_-treated plants of WT, *gln2* and *<u>glu1</u>* chl-roGFP2 expressing lines. DTT treatment resulted in a moderate decline in the 400/488 ratio compared to the resting state. A similar dynamic range (400/488nm under fully oxidized state divided by 400/488 under fully reduced state) of approximately 5.5 was recorded for all chl-roGFP2 lines. These results demonstrate the sensitivity of the probe to changing redox conditions and are consistent with probe characteristics previously observed in plant cells (Schwarzländer et al., 2008; Haber et al., 2021; Hipsch et al., 2021).

**Figure 4:**
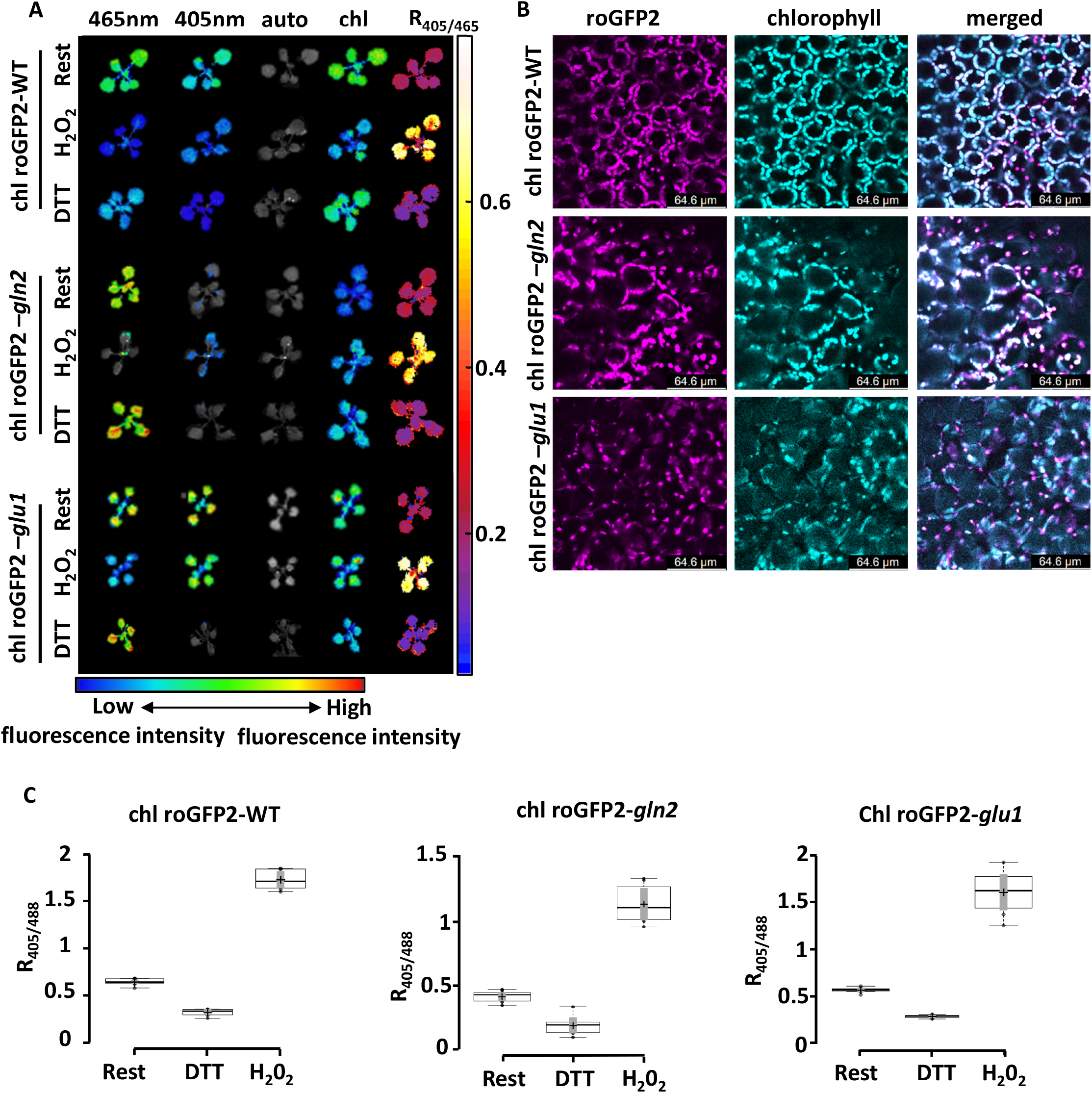
Redox sensitivity of chloroplast-targeted roGFP2 expressed in *gln2* and *glu* mutant lines. (A) Whole-plant images of chl-roGFP2 fluorescence following excitation at 405 nm or 465 nm (emission at 515 nm), autofluorescence following excitation at 405 (emission at 448nm), chlorophyll autofluorescence, and ratiometric analysis are shown for plants at rest and under fully oxidized (1000 mm H_2_O_2_, 10 min) and fully reduced [100 mm dithiothreitol (DTT), 60 min] conditions. Images were digitally combined for comparison. (B) Confocal images showing the fluorescence signals emanating from the three chl-roGFP2-expressing lines (chl-roGFP2 excitation: 488 nm). (C) Comparison between the roGFP2 ratio (400/485) at rest, under fully oxidized and fully reduced states, in WT and mutant lines. Fluorescence intensity in 3-week-old plants (n=12) was measured using a plate reader.

WT, *gln2* and *glu1* lines expressing chl-roGFP2 were exposed to 3 h of normal growth conditions (120 µmol m^−2^ s^−1^) at the beginning of the day, followed by a phase of 800 or 1200 µmol m^−2^ s^−1^ for a 10 h period. After the HL phase, the light was dimmed to the normal growth conditions for an additional 3 h period. chl-roGFP2 oxidation degree (OxD) was measured using an automated system that continuously measures fluorescence signals from living plants (Haber et al., 2021; Lampl et al., 2022). The chl-roGP2 OxD values were normalized with the OxD values collected under NL, measured the day before the experiment began. Upon transition from NL to HL conditions, an increase in chl-roGFP2 OxD was observed in all lines (Fig. 5A-B). Interestingly, when plants were returned to NL after the HL period, a decrease in chl-roGFP2 OxD to steady-state levels was clearly seen for WT but not for *gln2* and *glu1* lines. A return to steady state levels was only detected in the mutant lines when plants were placed in the dark (Fig. 5A-B).

**Figure 5:**
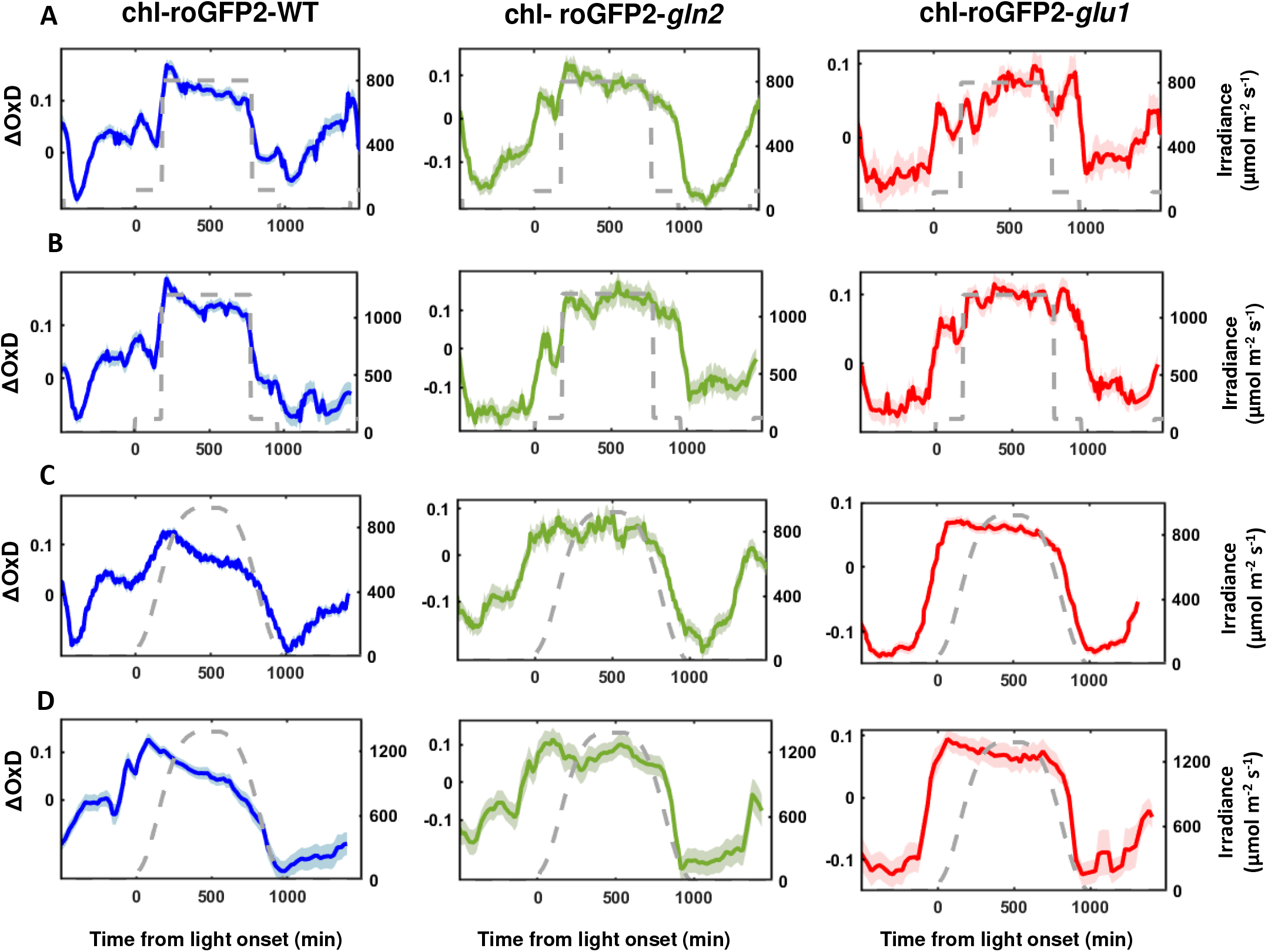
chl-*E*_*GSH*_ relaxation under high light involves nitrate assimilation activity. Diurnal changes in chl-roGFP2 degree of oxidation (OxD) under high light (HL) conditions in the *gln2* and *glu* mutant lines are shown. WT, *gln2* and *glu* lines were subjected to continuous HL conditions of 800 (A) or 1200 m^−2^ s^−1^ (B) or to gradually increasing light conditions peaking on 800 (C) or 1200 m^−2^ s^−1^ (D). The applied light intensities are depicted as a dashed line. Monitoring was performed in 3-week-old plants grown in 12-well plates. Each treatment involved 72-84 plants divided into 6-7 independent plates consolidated in a “sliding window” (n = 6-7) display. Values represent means, with SE presented as shaded regions.

When chl-roGFP2-expressing WT plant lines were exposed to gradual increases in light intensities, peaking at 800 or 1200µmol m^-2^s^-1^, followed by a gradual decrease, identical to the conditions introduced in the ΦPSII experiments (Fig. 5C-D), OxD gradually increased, peaking (ΔOxD= 0.127) when the light intensity was ∼600 µmol m^-2^s^-1^. From that point onward, a gradual decrease in chl-roGFP2 oxidation was observed, reaching its lowest oxidation degree (ΔOxD= - 0.113) at the beginning of the dark period. A similar pattern of initial oxidation followed by reduction during the HL period was also observed in plants exposed to 1200 µmol m^-2^s^-1^ (Fig. 5 C-D). These results agree with our previous reports showing oxidation followed by relaxation of chl-roGFP2 in repose to increasing light intensities (Lampl et al., 2022). Remarkably, chl-roGFP2 oxidation patterns in roGFP2-expressing *gln2* and *glu1* lines were different than those of WT. When exposed to increasing light intensities, mutant lines showed increased chl-roGFP2 oxidation, similar to WT, suggesting that the nitrate assimilation pathway does not affect the initial HL-driven changes in chl-*E*_*GSH*_. However, no relaxation was observed during the HL phase in the mutant lines. (Fig. 5C-D). Inspection of the chl-roGFP2 oxidation patterns in these lines revealed that the reduction in chl-roGFP2 OxD values occurred when light intensities were decreased towards darkness. These results demonstrate the effect of nitrogen assimilation in alleviating oxidative stress under HL and suggest that the reduction in chl-roGFP2 OxD during the HL period in WT plants is achieved by diverting excess electrons to the nitrogen assimilation pathway.

## Discussion

Tight coordination between the production of highly energized electrons in the photosynthetic light reaction and their consumption by downstream carbon reactions is crucial for preventing the overproduction of photosynthesis-generated ROS. Several photoprotective mechanisms have evolved in plants to protect the photosynthetic machinery from over-reduction of the photosynthetic electron chain. In some of them, energy is not recaptured (for example, quenched through heat production), while in others, energy is restored by rerouting electrons to alternative electron-consuming pathways, such as starch biosynthesis (Saroussi et al., 2019). While starch biosynthesis consumes electrons by being a repository for fixed carbon, nitrate assimilation captures electrons upstream to the CBC enzymes by consuming photosynthesis-derived reduced Fd and NADPH transported from the chloroplast through the malate shunt (Noctor and Foyer, 1998). Accordingly, it has been suggested that nitrogen assimilation plays a photoprotective role, avoiding energy imbalance induced by excessive reducing power and ATP accumulation (Noctor and Foyer, 1998; Busch et al., 2018). Using plants mutated in the nitrogen assimilation pathway, this study aimed to experimentally test the effectiveness of nitrogen assimilation in alleviating HL stress.

By tracing the incorporation of ^15^N-nitrate into amino acids, we demonstrated decreased nitrogen assimilation activity in *gln2* and *glu1* mutant lines under normal and HL conditions, confirming the role of GS2 and GOGA1 in the *de-novo* nitrate assimilation pathway. Yet, while lower than WT, the noticeable rate of nitrogen assimilation observed in these two mutant lines suggests that other molecular components can partially compensate for the lack of GS2 and GOGAT1. In the case of GS2, it was suggested that cytosolic GS activity could compensate for the lack of GS2 (Lam et al., 1996). Similarly, it is possible that GOGAT2 activity partially compensates for the lack of GOGAT1 activity in *glu1*, although GOGAT2 expression is mainly concentrated in roots (Coschigano et al., 1998). Alternatively, the activity of NADH-GOGAT may compensate for the lack of Fd-GOGAT1 activity (Lancien et al., 2002).

Besides being an electron consumer, nitrate assimilation also affects photosynthesis by its interconnection with the photorespiration pathway. This interaction arises from the dual role of the chloroplastic GS/GOGAT cycle in nitrate assimilation and photorespiration and the need for photorespiratory carbon skeletons for amino acid biosynthesis (Noctor and Foyer, 1998; Broncano et al., 2023). Indeed, inhibition of nitrate assimilation has been demonstrated under non-photorespiratory conditions, while improvement in carbon assimilation rates has been achieved upon dual activation of the nitrate assimilation and photorespiratory pathways (Rachmilevitch et al., 2004; Busch et al., 2018). The viable phenotype of *gln2* mutant line agreed with its recent characterization and suggests that it is not involved in the photorespiratory pathway (Hachiya et al., 2021; Lee et al., 2022). This was supported by the comparable maximum quantum efficiency of PSII in WT and *gln2* (Fig. 4), in contrast to the enhanced levels of photoinhibition reported in photorespiration-defective mutant lines upon exposure to air (Takahashi et al., 2007). The typical photorespiratory phenotype of *glu1*, which developed into small, chlorotic plants with low maximum quantum efficiency of PSII when grown under air, was in line with previous observations (Somerville and Ogren, 1980; Coschigano et al., 1998). The low incorporation of ^15^N into Gln and Glu in this line (Fig. 3C) confirmed the role of Fd-GOGAT in the de-novo nitrate assimilation pathway, positioning Fd-GOGAT in a central position in plant nitrogen metabolism. Due to its dual function in photorespiration and nitrate assimilation, the observed changes in ΦPSII and chl-*E*_*GSH*_ under HL in *glu1* cannot be exclusively attributed to de-novo nitrogen assimilation pathways.

The pronounced decrease in ΦPSII in both mutants under HL (Fig. 3) highlights the importance of nitrate assimilation in electron consumption to alleviate PSII photodamage. In line with these observations, increased activities of NR and GS were recently reported in cotton plants exposed to HL conditions. In addition, plants with high nitrate supply showed favorable photochemical characteristics such as higher electron flux through PSII and lower limitation in the acceptor side of PSI than those provided with minimal nitrate levels (Guilherme et al., 2019). Considering the high sensitivity of PSI to conditions in which electron flow from PSII exceeds the capacity of their consumption by PSI electron acceptors (Sonoike, 2011), the higher PSII photoinhibition levels observed in the mutant lines can be viewed as an alternative photoprotection mechanism (Tikkanen et al., 2014) that is enhanced when the diverting of reducing power to nitrate assimilation is not adequate to prevent overreduction of the photosynthetic electron transfer chain. This hierarchy may indicate the higher tendency to route incoming energy to sustainable sinks rather than quenching through heat production. The importance of nitrate assimilation in managing excitation energy downstream to PSI was evident by the observed chl-*E*_*GSH*_ patterns. Specifically, chl-roGFP2 relaxation, observed upon transfer of WT plants from HL to NL, was not observed in the mutant lines. In addition, chl-roGFP2 oxidation was reported in nitrogen-starved diatom cells (Rosenwasser et al., 2014b). chl-roGFP2 oxidation state reflects the balance between H_2_O_2_ production and NADPH availability for GR activity (Meyer et al., 2007; Meyer, 2008). As nitrate assimilation does not directly contribute to GR activity, these results suggest that the diversion of electrons for nitrate assimilation reduces ROS production, contributing to the regulation of the chl-*E*_*GSH*_ homeostasis under HL.

As a major electron-consuming pathway, nitrate assimilation activity is tightly linked to the photosynthetic light and carbon reactions. The tight regulation of electron partitioning between nitrate assimilation and other electron-consuming pathways is achieved, at least in part, by the redox regulatory systems, as indicated by the redox sensitivity and Trx-based regulation of various N assimilation enzymes (Lemaire et al., 2007; Rosenwasser et al., 2014; González et al., 2019). The effect of nitrate assimilation on the chloroplast redox homeostasis and the sensitivity of nitrogen assimilation enzymes to redox modifications may enable feedback mechanisms in which the nitrate assimilation rates are continuously adjusted according to ROS levels and NADPH availability.

The compromised nitrate assimilation observed in the two mutant lines may affect multiple cellular processes involving nitrogenous compounds. This may make it challenging to discern between the effect of the nitrate assimilation per se and the potential effects of inadequate nitrogen availability. For example, alterations in chloroplastic *E*_*GSH*_ may be a result of the decreased GSH content, as the synthesis of amino acids might be limited as nitrate assimilation is less efficient. Yet, several lines of evidence suggest a direct link between nitrate assimilation, photosynthetic efficiency and GSH redox state. The similar ΦPSII levels detected in WT and *gln2* under normal growth conditions (Fig. 3) indicate that the vulnerability of *gln2* to HL is directly caused by impairments in nitrogen assimilation pathways rather than a global change in leaf nitrogen status. In contrast, the low ΦPSII levels in *glu1* were likely related to the overall defects of this mutant when grown in ambient CO_2_ levels. Indeed, extensive changes in gene expression profiles in *glu1* plants have been demonstrated in transcriptomic analyses, including the downregulation of photosynthesis-related pathways and the induction of stress-related genes (Kissen et al., 2010). Notably, despite the poor phenotype of *glu1*, chl-roGFP2 OxD values were similar in WT and *glu1*, under steady-state and in response to H_2_O_2_, suggesting no differences in overall antioxidant levels between the plant lines (Fig. S5). Importantly, the similar deviation from WT patterns observed for chl-roGFP2 OxD in *gln2* and *glu*, despite the apparent differences in their phenotypes (Fig. 5), strengthens the notion that de-novo nitrate assimilation serves as an electron sink, which contributes to the maintenance of redox homeostasis under HL.

Nitrate and ammonium are the primary nitrogen forms absorbed by plant roots, and their relative availability varies in different ecosystems and environmental conditions (Reed and Hageman, 1980; Andrews, 1986). Under light-limiting conditions, ammonia assimilation can be an adaptive strategy to fulfill the heightened demand for photosynthetically derived reducing power when nitrate is assimilated (Debiasi et al., 2021). Yet, ammonium might be toxic to plant cells when it accumulates to high concentrations, limiting its utilization even under low light conditions. On the other hand, in environments where light is not a limiting factor and especially in stressful fluctuating environments where the risk for photodamage is high, the higher energetic requirements of nitrate compared to ammonium assimilation, renders nitrate assimilation a stronger sink for electrons and, therefore a better strategy to cope with HL-induced stress. Indeed, faster net oxygen evolution was found in plants exposed to high light when receiving nitrate rather than ammonium as a nitrogen source (Walker et al., 2014; Bloom, 2015). Therefore, sophisticated nitrogen fertilizer management, according to the spatiotemporal variability in light conditions in agricultural fields, could decrease PSII photodamage and slow NPQ activation and likely improve photosynthesis and crop yields. Additional experiments will be needed to determine if nitrogen fertilization protocols can be fine-tuned to both provide plants with adequate nitrogen and to allow plants to better acclimate to HL-induced stress conditions.

Nitrate assimilation occurs both in roots and shoots, with high variation across plant species in their reliance on such activities in the root versus shoot, as explored by the differences in NR activity in the two plant parts and the relative abundance of nitrate and reduced nitrogen compounds in the xylem sap of various plant species (Crafts-Brandner and Harper, 1982; Sprent and Thomas, 1984; Smirnoff and Stewart, 1985; Andrews, 1986; Debiasi et al., 2021). Root assimilation, which relies on respiration-derived reductants, is more energetically costly than utilizing reducing equivalents directly from the photosynthetic light reaction, although the reliance on respiratory fixed sugar was suggested to be an advantage in certain conditions, such as low temperatures (Andrews, 1986). While the mechanisms that determine the partitioning of nitrate assimilation between root and shoot have not been resolved, the presented role of nitrate assimilation in protecting PSII photodamage and regulating the chloroplastic GSH redox state might be a key factor in adopting root versus shoot assimilation strategies.

## Methods

### Plant material and growth conditions

*Arabidopsis thaliana* WT (ecotype Columbia-0), *gln2* (SALK_051953, At5g35630) and *glu1* (CS8611, At5g04140) lines were used throughout this research. Seeds were sown on unfertilized Klasmann-Deilmann (K®) Select soil (fertilized manually at sowing with 1g/L of 20-20-20+Micro [E5020372KJSS12, Ecogan]), transferred to the dark at 4 °C for two days and then grown under a 16/8 light-dark cycle with a photosynthetic photon flux density (PPFD) of 120 μmol m−2 s−1 (21ºC, 60-70% RH, ambient CO_2_), for 2-3 weeks. For ^15^N incorporation experiments, plants were grown in hydroponic nutrient solution for 3 weeks.

### Hydroponics solution

The hydroponics nutrient solution, based on Hoagland solution, comprised of 2M KNO_3_ (or K^15^NO_3_), 1M Ca(NO_3_)_2_, 1M MgSO_4_, 1M KH_2_PO_4_ and 0.1M EDFS as macroelements and 0.5mM CuSO_4_, 0.5mM H_2_MoO_4_, 2mM MnSO_4_, 50mM KCl, 25mM H_3_BO_3_ and 2mM ZnSO_4_ as microelements. The solution pH was set to 5.7-5.8 using NaOH and HCl for adjustments (measured by Jenway 351000 3510 Bench pH/mV Meter).

### LC-HRMS analysis of glutamine and glutamic acid

Frozen plant samples (5-60 mg, depending on plant line) were homogenized in a Bioprep homogenizer using 2 mm zirconium oxide beads. Amino acids were extracted from homogenized samples with 1 ml acetonitrile/water (with 20 mM ammonium formate) (1:1) solution following centrifugation (at 10,000 rpm for 10 min) and filtration of supernatant through membrane filters (regenerated cellulose, Teknokroma, Spain). Samples were analyzed by a LC-MS system, which consisted of a Dionex Ultimate 3000 RS HPLC coupled to a Q Exactive Plus hybrid FT mass spectrometer equipped with a heated electrospray ionization source (Thermo Fisher Scientific Inc.). The chromatographic separations of free amino acids were carried out using a HILIC-Z column (2.1×150 mm, particle size 2.7 µm, Agilent) employing a linear binary gradient of (A) acetonitrile and (B) water with 100 mM ammonium formate. The mass spectrometer was operated in positive ionization mode, with the following ion source parameters: spray voltage 3.5 kV, capillary temperature 250□C, sheath gas rate (arb) 40, and auxiliary gas rate (arb) 10. Mass spectra were acquired in the m/z 70-300 Da range at resolving power 140.000. The highest possible resolving power was used for splitting naturally occurring 13C isotope from experimentally introduced 15N isotopes. The LC-MS system was controlled, and data were analyzed using the Xcalibur software (Thermo Fisher Scientific Inc.). The relative amount of ^15^N_1_-Gln was calculated using the following equation:

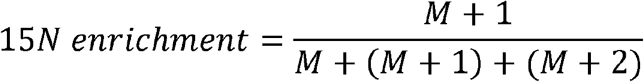

For the quantification of labeled ^15^N_2_-Gln, the above equation was used with M+2 in the numerator.

### Western blotting

Protein extracts were prepared, using a hand homogenizer, in 2 ml tubes with 200 µL extraction buffer containing 5µL/mL NP40 and 1µL/mL protease inhibitor (PI). Proteins were precipitated for 2 min on ice and centrifuged at 14,000 rpm for 17 min, at 4 ºC. The supernatant (100µL) of each sample was transferred to fresh 2 ml tubes. DTT (10 mM, final concentration) and sample buffer (x3) (150 mM Tris-HCl, pH 6.8, 6% (w/v) SDS, 30% glycerol and 0.3% pyronin Y) were added to each sample. The samples were then boiled for 10 min and immediately transferred to ice for 2 min. Gel fractionation was carried out on handcast 10% polyacrylamide gel. Fractionated proteins were transferred to a polyvinylidene fluoride membrane (Bio-Rad), using the Trans-Blot Turbo Transfer System (Bio-Rad) with Trans-Blot Turbo Midi Transfer Packs. The membrane was incubated with anti-GS2 antibody (1:5000; agrisera) or anti-GOGAT1 antibody (1:1000; agrisera), both followed by goat anti-rabbit horseradish peroxidase (HRP)-conjugated IgG (1:20,000; agrisera).

Chemiluminescence was detected using the Advanced Molecular Imager HT (Spectral Ami-HT, Spectral Instruments Imaging, LLC., USA).

### chl-roGFP2 transformation

The entire gene sequence of chl-roGFP2 was chemically synthesized for the generation of a chl-roGFP2-expressing line. Chloroplast targeting was achieved using the 2-Cys peroxiredoxin A (PRXa; Uniprot ID: Q96291) signal peptide. The chl-roGFP2 gene was cloned into the plant cloning vector pART7 using XhoI and HindIII restriction enzymes. The entire construct, including the CaMV 35S promoter and ocs terminator, was cloned into the binary vector pART27 using the restriction enzyme NotI. The pART27 plasmid containing the chl-PRXaTP-roGFP2 construct was transformed into Agrobacterium tumefaciens (GV3101) using heat-shock transformation.

The transformed Agrobacterium were grown on selective plates (LB agar [35g/1L dW] + rifampicin [10 ng/ml], gentamicin [30 ng/ml] and spectinomycin [100 ng/ml]), at 28 °C for 24-72 h (until visible bacterial coverage). The bacteria were then re-suspended in 50 ml falcons containing 20 ml LB, supplemented with rifampicin (10 ng/ml), gentamicin (30 ng/ml) and spectinomycin (100 ng/ml) and then incubated (28 °C, shaken at 200 rpm) until OD=0.8-1 (measured using a MRC V-1100D spectrophotometer). The tubes were then centrifuged for 5 min (4 °C, 4000 rpm) and pellets were re-suspended in 10 ml 1% sucrose in distilled water (dW) to OD=1, after which, 0.03% 77-L (Agan Adama) was added (referred to below as Transformation Solution). Transformation of *A. thaliana* was performed by floral dip (Clough and Bent, 1998). Transformed lines were selected based on kanamycin resistance and the chl-roGFP2 fluorescence signal.

### Confocal microscopy

Images were acquired with a Leica TCS SP8 confocal system (Leica Microsystems) using the LAS X Life Science Software and an HC PL APO ×40/1.10 objective. All images were acquired at 4096 × 4096 pixel resolution, with an emission wavelength of 500-520 nm and excitation of 488 nm for chl-roGFP2 fluorescence and emission wavelength of 670 nm and excitation of 488 nm for chlorophyll fluorescence. All images were generated using Fiji (Image J) software.

### Chl-roGFP2 fluorescence measurements and analysis

Whole plant redox imaging was performed on 10 days-old transgenic chl-roGFP2-expressing plants. Fluorescence was detected using an Advanced Molecular Imager HT, and images were taken using the AMIview software. Plants were excited with 405 nm±10 or 465 nm±10 LED light sources and fluorescence was measured through a 515 nm±10 emission filter. For chlorophyll detection, a 405 nm±10 LED light source and a 670 nm±10 emission filter were used. All images were captured under the following settings: exposure time = 1 s, pixel binning = 2, field view (FOV) = 25 cm, and LED excitation power 40% and 60%, for 405 nm and 465 nm excitations, respectively. Excitation power for chlorophyll detection was 5%. Chlorophyll autofluorescence was measured to generate a chlorophyll mask, which was then used to select pixels that returned a positive chlorophyll fluorescence signal. Only those pixels were subsequently considered for the roGFP analysis. For autofluorescence correction, excitation was performed with a 405 nmL±L10 LED and a 448 nmL±L10 emission filter was used; the excitation power was 60%. Autofluorescence-corrected ration metric images were generated according to Hipsc et al. (2022)

Quantitative chl-roGFP2 analysis was mainly carried out as per the plate reader method according to (Haber et al., 2021; Lampl et al., 2022) using the Spark® Multimode Microplate Reader (Tecan, Switzerland). The following filters were used: excitation: 400/20 nm and 485/20 nm; emission: 520/10 nm. For chlorophyll detection, excitation at 400/20 nm and emission at 670/40 nm. For automatic detection of chl-roGFP2 signals, plants growing in 12-well plates were placed in a Fytoscope FS-SI-4600 (PSI, Czech Republic) chamber and were automatically taken into the fluorometer for fluorescence detection using a KiNEDx KX.01467 robot (paa, UK and USA). A 9-by-9-pixel matrix was formed from each well. Chlorophyll fluorescence was detected at 400 nm/670 nm to create a chlorophyll mask. Only pixels with a positive chlorophyll fluorescence signal were taken into account for the roGFP analysis. Then, the average fluorescence values recorded in non-fluorescent plants (WT) were calculated, and the values were subtracted as background autofluorescence from the values measured in the chl-roGFP2 plants. roGFP2 degree of oxidation (the relative quantity of oxidized roGFP proteins, OxD) was calculated based on the fluorescence signal, as previously described (Meyer et al., 2007).

### Chlorophyll fluorescence measurements

Chlorophyll fluorescence was measured using a Walz PAM IMAGING PAM M-series IMAG-K7 (MAXI) fluorometer. The effective PS II quantum yield (ΦPSII) was determined by the equation of ΦPSII=(FmL-F)/FmL. Images were analyzed with ImagingWinGigE V2.56p software.

### Statistics

For ^15^N enrichment analysis, statistical significance was tested using a two-tail Student’s t test using JMP, and is indicated by asterisks.

