## Supplementary Data for "Nitrogen assimilation plays a role in balancing the chloroplastic glutathione redox state under high-light conditions"

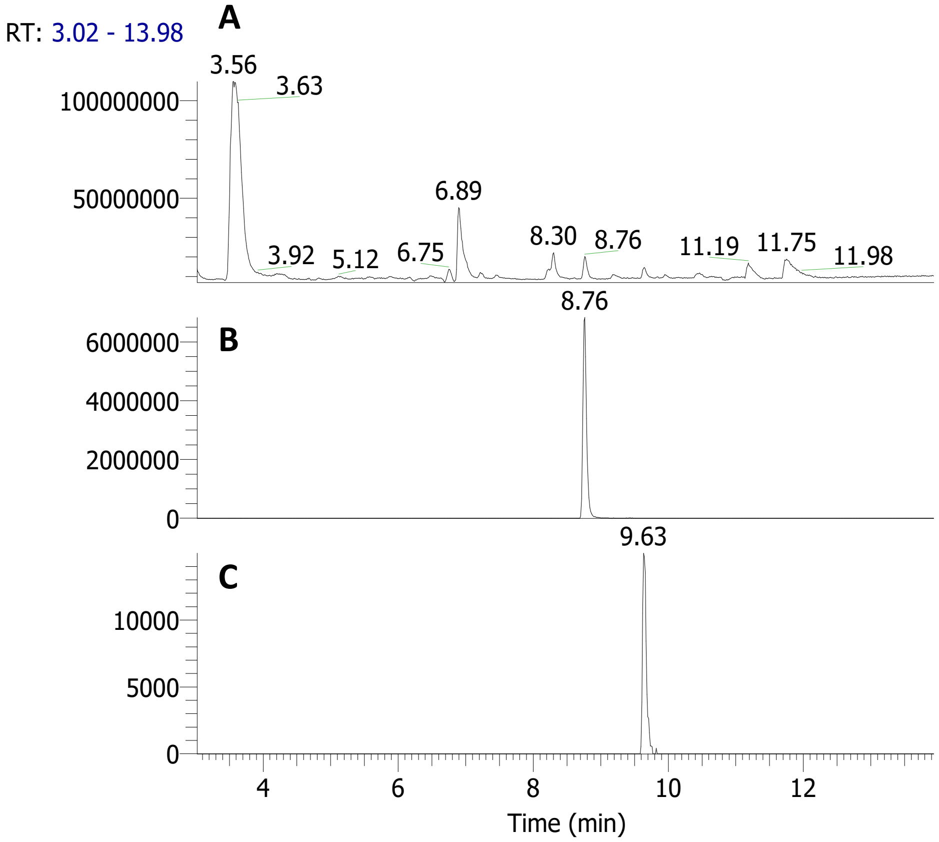


**Figure S1:** An example of positive ESI LC-MS chromatograms of leaf extract. (A) TIC (total ion count) chromatogram. (B&C) EIC (extracted ion count) chromatograms of glutamine (B, RT 8.76 min) and glutamic acid (C, RT 9.6 min).

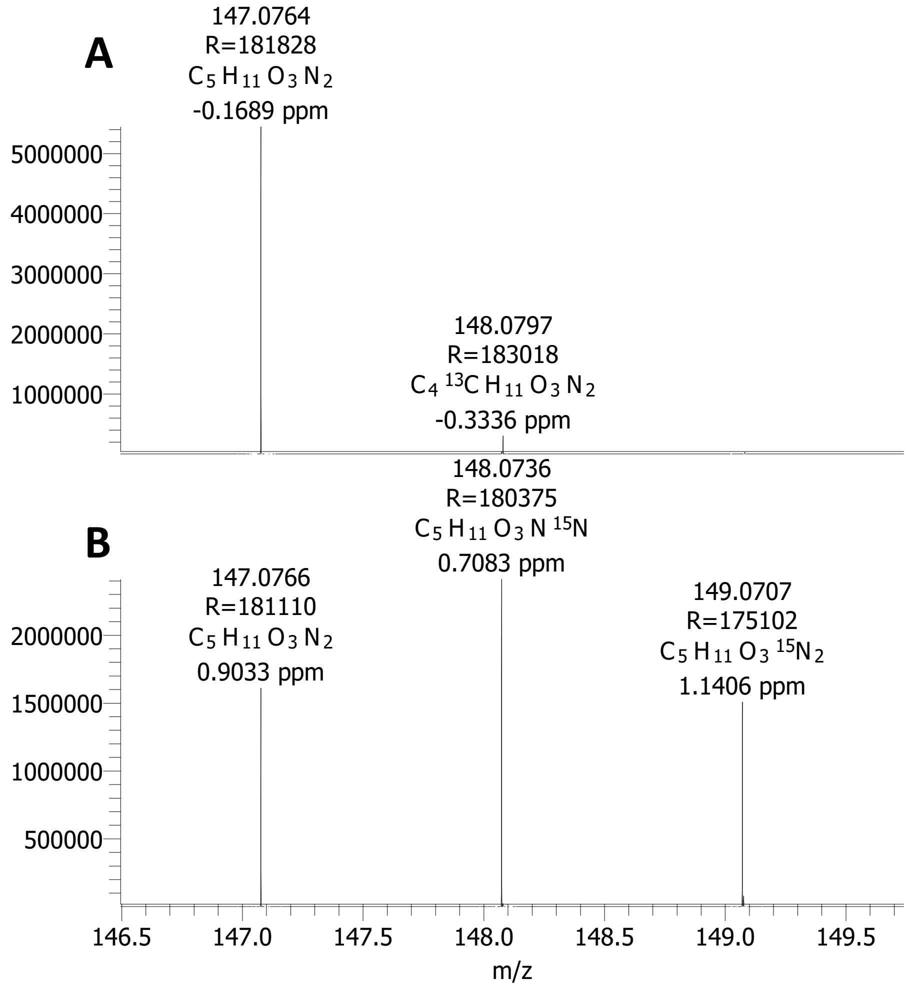


**Figure S2: Descerning between 15N and 13C isotopes detected in M+1 ion of glutamine**. High resolution (resolving power >180,000) positive ESI mass spectrum of glutamine detected in unlabeled sample (A) and sample labeled with $^{15}N$ nitrate

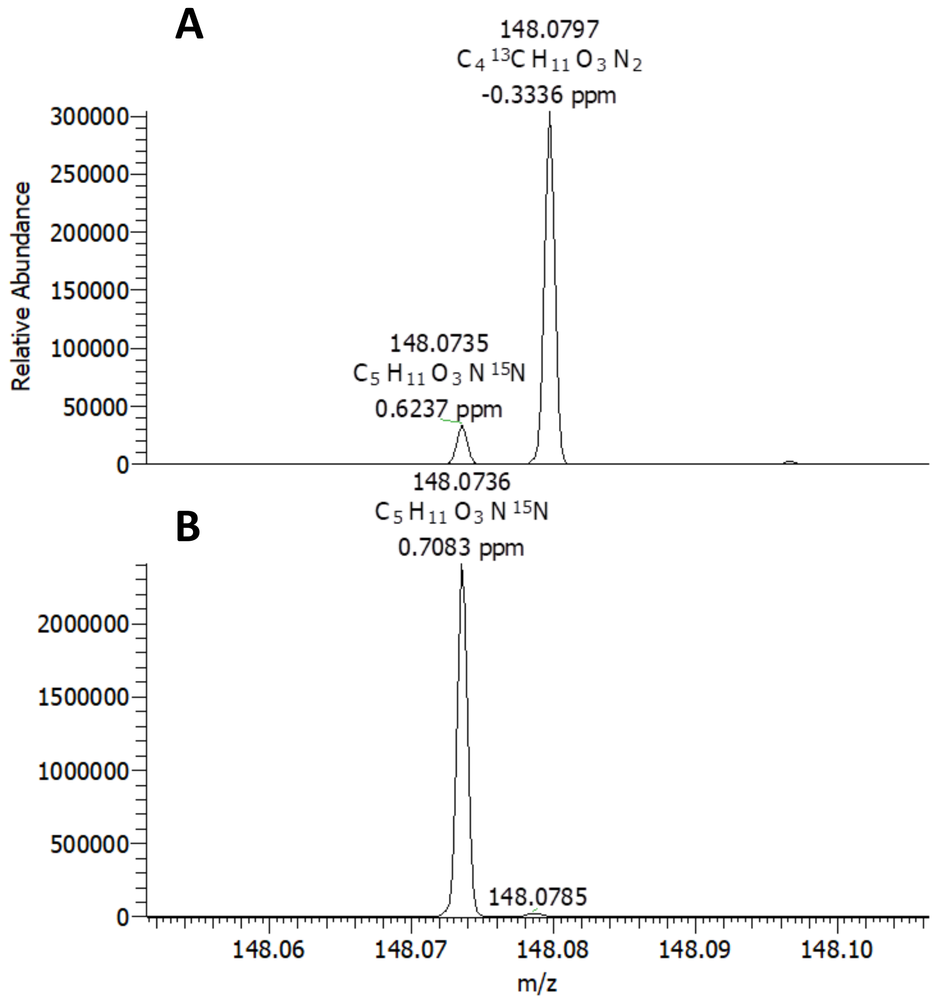


**Figure S3: Complete resolution of $^{15}N$ and $^{13}C$ isotopes detected in M+1 ion of glutamine.** Chromogram showing the relative abundance of $^{15}N$ Gln (left peak), and $^{14}N$ Gln containing $^{13}C$ (right peak) in unlabeled sample (A) and $^{15}N$-labeled sample (B).

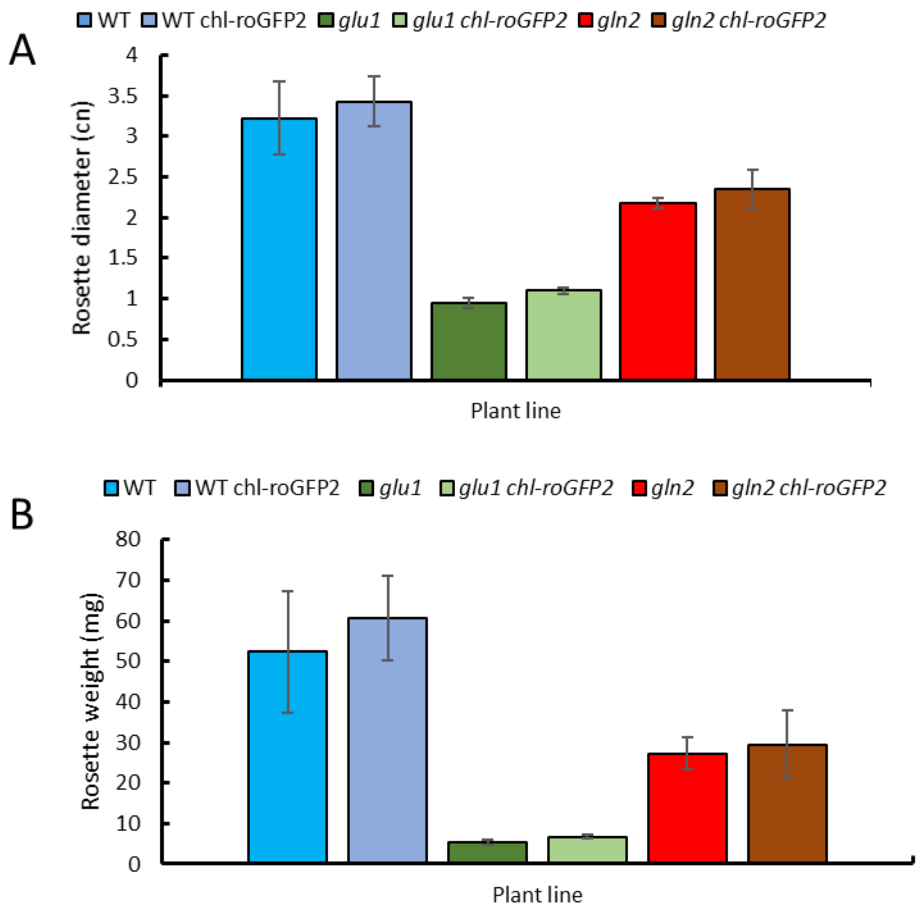


**Figure S4: A comparison between the phenotypes of each Arabidopsis thaliana line to their correspondence chl-roGFP2 insertion line**. Rosette diameter (A) Rosette weight (B) at three weeks old are presented. Data shown for plants lacking chl-roGFP2 is equivalent to the data presented in Fig. 1.

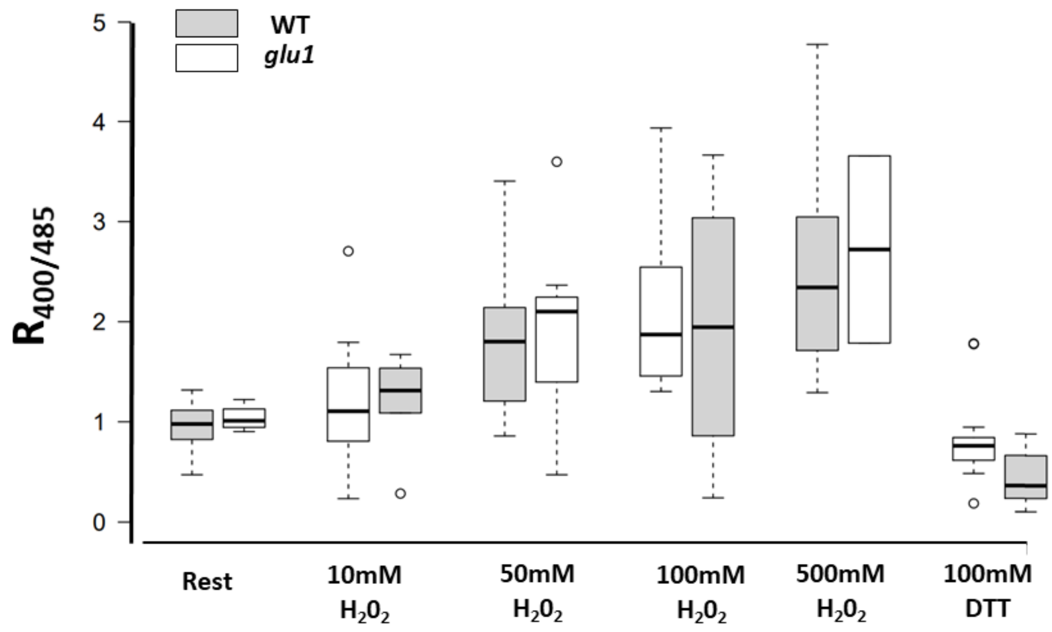


**Figure S5: The response of chl-roGFP2 to hydrogen peroxide in WT and *glu1* lines.** Flourescense ratio values (400/485) recorded in WT and *glu1* lines exposed to various hydrogen peroxide concentrations for 3 min or 60 min after DTT treatment are shown.
